# Comparison of *de novo* and reference genome-based transcriptome assembly pipelines for differential expression analysis of RNA sequencing data

**DOI:** 10.1101/2022.08.20.504634

**Authors:** Rebekah A. Oomen, Halvor Knutsen, Esben M. Olsen, Sissel Jentoft, Nils Chr. Stenseth, Jeffrey A. Hutchings

## Abstract

**Objective:** As sequencing technologies become more accessible and bioinformatic tools improve, genomic resources are increasingly available for non-model species. Using a draft genome to guide transcriptome assembly from RNA sequencing data, rather than performing assembly *de novo*, affects downstream analyses. Yet, direct comparisons of these approaches are rare. Here, we compare the results of the standard *de novo* assembly pipeline (‘Trinity’) and two reference genome-based pipelines (‘Tuxedo’ and the ‘new Tuxedo’) for differential expression and gene ontology enrichment analysis of a companion study on Atlantic cod (*Gadus morhua*).

**Results:** The new Tuxedo pipeline produced a higher quality assembly than the Tuxedo suite. However, greater enrichment of Trinity-identified differentially expressed genes suggests that a higher proportion of them represent biologically meaningful differences in transcription, as opposed to transcriptional noise or false positives. Coupled with the ability to annotate novel loci, the increased sensitivity of the Trinity pipeline might make it preferable over the reference genome-based approaches for studies aimed at broadly characterizing variation in the magnitude of expression differences and biological processes. However, the ‘new Tuxedo’ pipeline might be appropriate when a more conservative approach is warranted, such as for the identification of candidate genes.

## Introduction

RNA sequencing (RNA-seq) revolutionized ecological genomics by enabling transcriptome assembly without *a priori* genomic information [1, 2]. As sequencing technologies become more accessible and bioinformatic tools rapidly improve, genomic resources are increasingly available for non-model species [3]. The choice of whether to use a draft genome to guide transcriptome assembly or perform assembly *de novo* is one that is best addressed early on in an RNA sequencing experiment and has the potential to substantially influence the results obtained from differential expression (DE) and enrichment analyses. Yet, studies are, by and large, reporting the outcome of a single analytical pipeline that produced perhaps the most comprehensible result. This is exacerbated by the fact that length restrictions on journal articles, and even supplementary materials, encourage unhelpful reductions in the reporting of technical details and alternative methods that had been evaluated during the course of data analysis. Here, we analyze RNA-seq data from a companion study on a widely distributed marine fish that exemplifies the increasingly common position of straddling the realms of model and non-model species, the Atlantic cod (*Gadus morhua*) (Oomen et al. provisionally accepted at *Integrative & Comparative Biology*). Specifically, we compare *de novo* transcriptome assembly and two reference genome-based pipelines in the context of differential expression and gene ontology enrichment analysis in a larval rearing experiment. Our study will provide a useful resource for anyone planning or conducting transcriptomic analysis of non-model or semi-model species and will shed light on the issue of restrictive publication lengths in an age of rapidly expanding genomic tools and analytical pipelines.

## Main text

### Methods

#### Experimental design

Briefly, we raised laboratory-hatched larvae of wild origin at three temperatures (9°C, 11°C, and 13°C). We sampled a total of three larvae from two tank replicates at each temperature at 2, 14, and 29 days post hatch (dph), and an additional three larvae from the hatching tank at 0 dph (prior to transfer to temperature treatments) to serve as a baseline sample (n=30). For details see Oomen et al. (provisionally accepted at *Integrative & Comparative Biology*).

#### RNA library preparation and quality control

Total RNA was isolated from individual whole larvae, using the RNeasy Mini Kit (Qiagen) according to the manufacturer’s instructions. Tissue homogenization was carried out using a Fastprep-24 Instrument (MP Biomedicals) for 30 s at 4.0 M/S in 1.5 ml tubes containing ceramic beads and 250 μl 1x lysis buffer. RNA was eluted in two steps, using 25 μl of RNase-free water each. Quality of RNA isolates was evaluated with an Agilent 2100 Bioanalyzer (BioRad) before and after library preparation with the TruSeq™ RNA low-throughput protocol (Illumina) and a fragmentation time of 4 minutes. Trimming and adapter removal was performed on all sequences, using Cutadapt v.1.8 [4] with a quality threshold of 20, followed by a hard trim of 10 from the 5’ end with a minimum remaining sequence length of 20 (Table 1). Sequence quality was evaluated before and after each trimming step, using FastQC v.0.11.2 [5]. Transcriptome assembly was performed with the remaining 740 million read pairs. Default parameters were used unless otherwise stated.

**Table 1:**
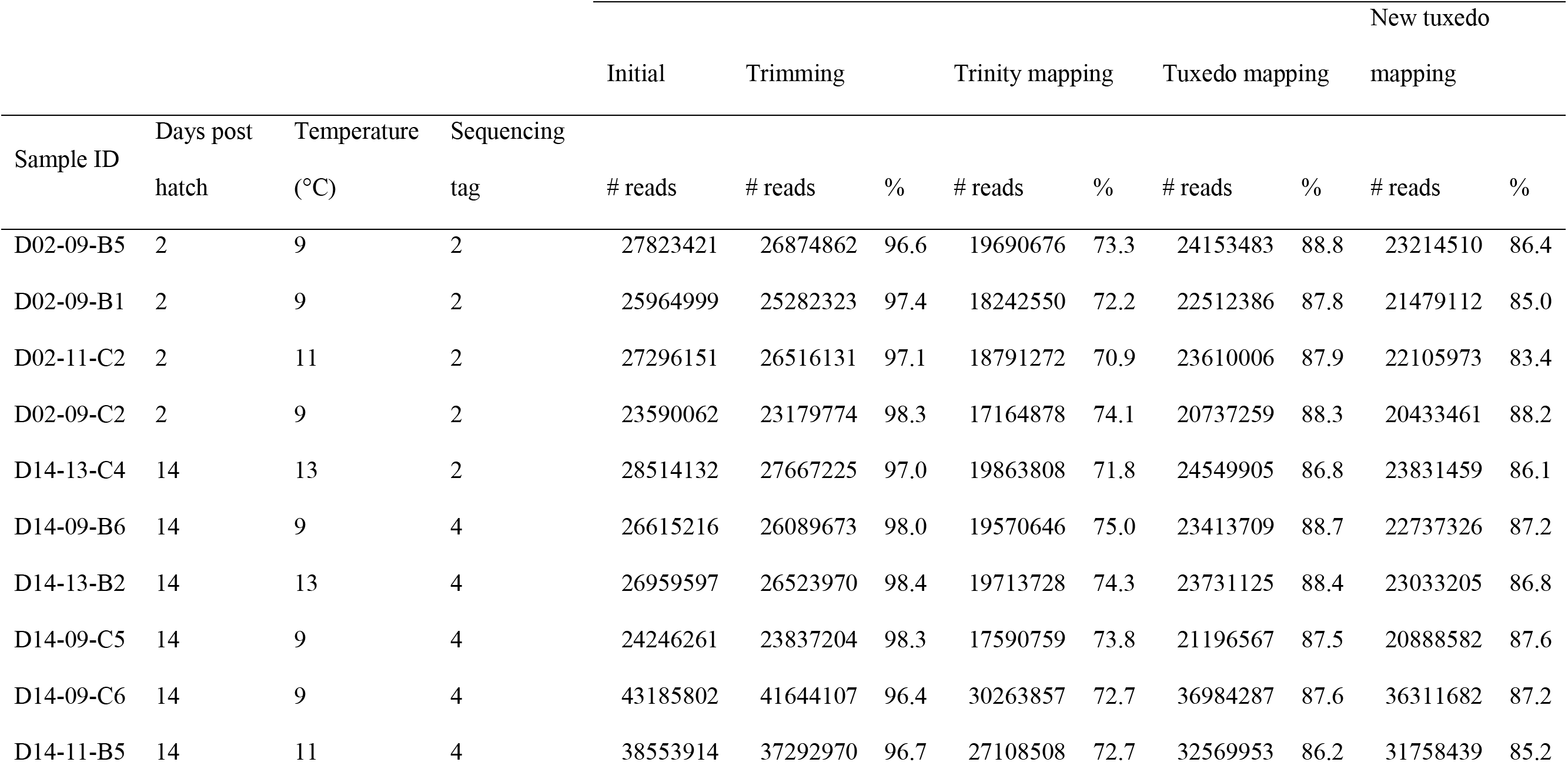
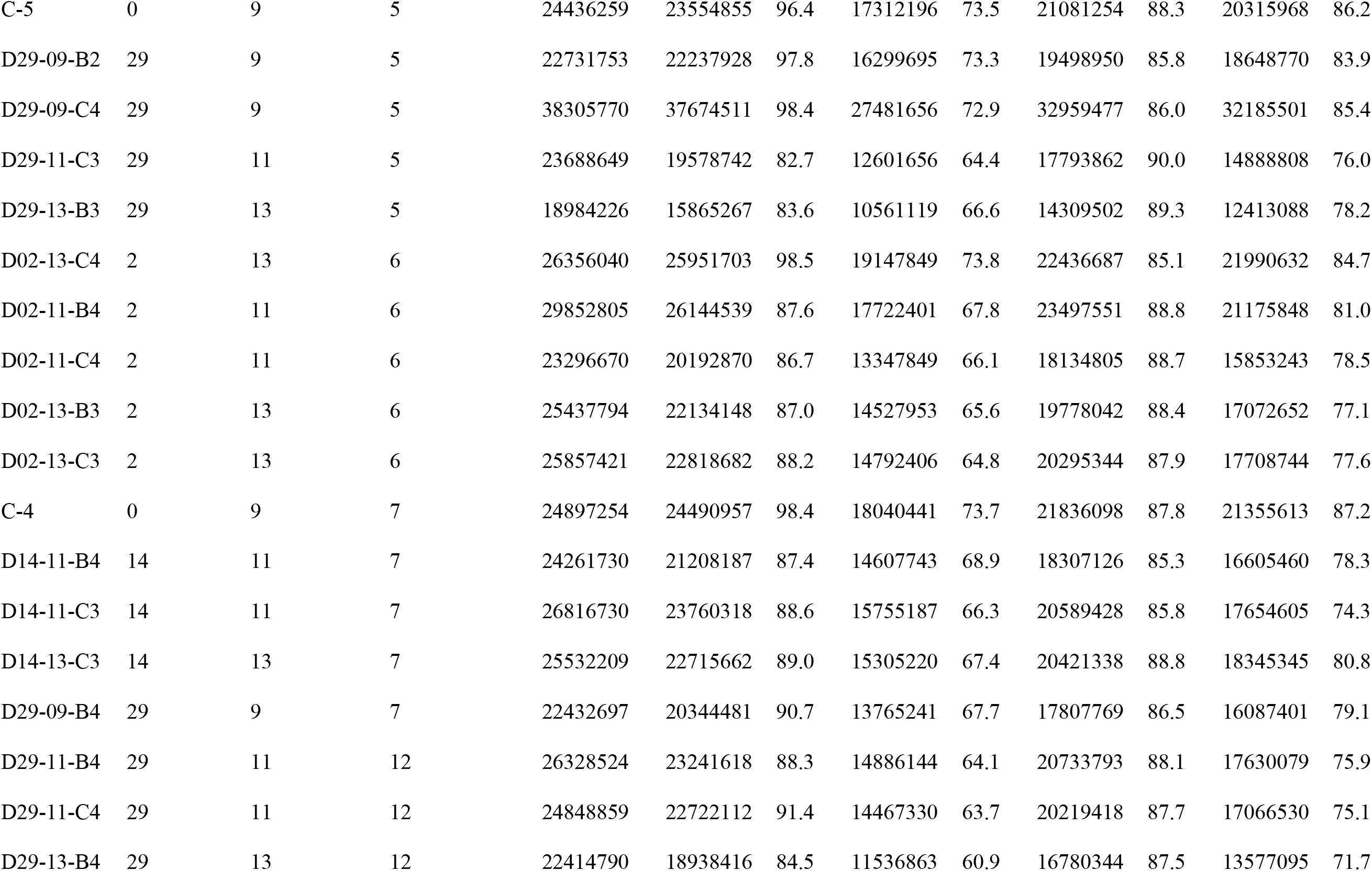
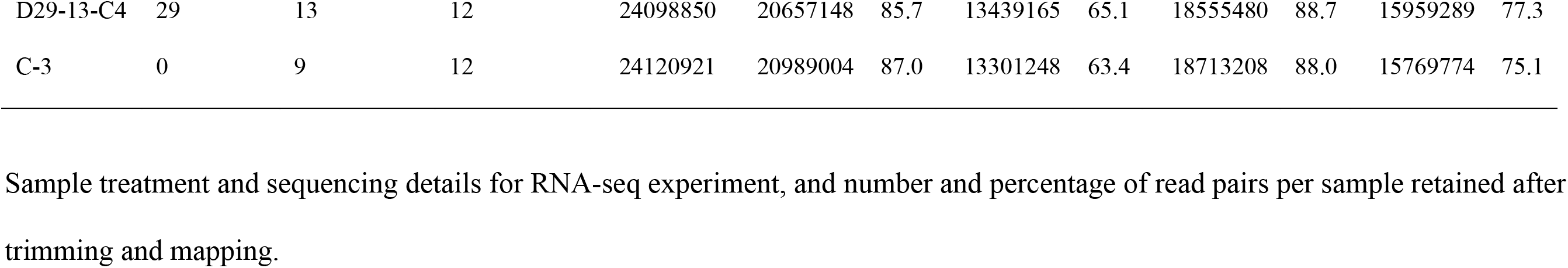
Sample information.

#### De novo *transcriptome assembly*

The transcriptome was assembled *de novo*, using the Trinity software suite v.2014-07-17 (http://trinityrnaseq.github.io), including bowtie v.1.0.0 [6] and samtools v.0.1.19 [7]. Trinity was performed with built-in normalization, 50 GB maximum memory and 10 CPUs. Assembly statistics were generated with TrinityStats.pl. The assembly was further evaluated for quality by calculating the contig ExN50 statistic, using contig_ExN50_statistic.pl, evaluating the relationship between raw read counts and FPKM (Fragments Per Kilobase of transcript per Million mapped reads), using count_features_given_MIN_FPKM_threshold.pl, and by counting the number of full-length transcripts using blastx and blastp (https://blast.ncbi.nlm.nih.gov/Blast.cgi) with the analyze_blastPlus_topHit_coverage.pl script. Reads were mapped back to the assembly, using align_and_estimate_abundance.pl with RSEM estimation and the bowtie aligner, to generate sample-specific transcript counts. The assembly was annotated using Trinotate v.2.0.1 (http://trinotate.github.io/).

#### Reference genome-based transcriptome assembly

In the Tuxedo pipeline, sequences were mapped to the second version of the cod genome [8], using Tophat v. 2.1.1 [9] with a mate inner distance of 200±200 bp. Cufflinks v.2.2.1 [10] was used to assemble sample-specific transcriptomes in reference-annotation-based transcript assembly mode [11] with fragment bias correction [12], which were then combined into a single assembly with Cuffmerge. Global pairwise DE analysis was performed using Cuffdiff and cummeRbund v.2.8.2 in R v.3.3.2.

In the ‘new Tuxedo’ pipeline, the low-memory aligner HISAT v.2.1.0 [13] mapped reads to the reference genome [8]. Samtools v.1.3.1 [7] sorted and converted the resulting SAM files to BAM files for transcript assembly, quantification, and merging of assemblies, using StringTie 1.3.1 [14]. We included a reference annotation containing known gene models in StringTie to improve reconstruction of low-abundance genes. Otherwise, default options were used. A matrix of raw gene counts was generated from the StringTie output, using the prepDE.py script (http://ccb.jhu.edu/software/stringtie/dl/prepDE.py). The assemblies resulting from both reference-based pipelines were evaluated relative to the reference genome and to each other, using gffcompare (https://github.com/gpertea/gffcompare).

#### Differential expression and gene ontology enrichment analysis

Differential expression analysis was conducted in edgeR v.3.16.5 [15] and included only those transcripts having a CPM (counts per million) >1 in at least three samples. We conducted pairwise comparisons between all samples compared to the baseline sample, using a false discovery rate (FDR)-corrected P-value of 0.05. Venn diagrams were constructed, using BioVenn [16] and Venny (http://bioinfogp.cnb.csic.es/tools/venny/). to compare differentially expressed gene lists.

Gene ontology (GO) enrichment analysis was performed, using ClueGO v.2.3.2 [17] and BiNGO v.3.0.3 [18] in Cytoscape v.3.2.1 [19], to identify significantly enriched biological processes. Up- and down-regulated genes were analyzed separately with an FDR-corrected P-value of 0.05 and otherwise default parameters. The Gene Fusion option was used in ClueGO.

### Results

#### Assembly

The *de novo* pipeline assembled 386,872 trinity ‘genes’ (i.e., unique groups of isoforms) corresponding to 530,683 unique transcripts (Additional file 1 Table S1). The peak N50 value of 2153 occurring at E90 suggests that sufficient sequencing depth was achieved and that approximately 41,413 biologically relevant transcripts were assembled (Additional file 1 Figure S1, Table S2). This roughly corresponds with the number of transcripts with an expression level >1 FPKM (Fragments Per Kilobase Million) (39,767; Additional file 1 Figure S2). Full-length transcript analysis, using blastx and blastp, detected 26,790 and 25,249 proteins, respectively, of which 9430 and 9774 were nearly full-length (≥90%) (Additional file 1 Table S3). Overall mapping rates of 61-75% resulted in approximately 10-30 million reads (mean = 17 million) per sample being successfully mapped back to the transcriptome (Table 1), meeting or exceeding coverage guidelines for DE analysis in a complex eukaryote [20].

The ‘new Tuxedo’ aligner HISAT performed approximately 10x faster and used much less memory than the Tuxedo suite aligner Tophat (data not shown). StringTie assembled slightly more loci (34,132) compared to Cufflinks (31,830) and approximately half (55%) as many unique transcripts, with generally higher precision (i.e., a lower apparent false-positive rate), suggesting a higher quality assembly with fewer misassembled transcripts (Additional file 1 Table S4). For both assemblies, approximately one-third of all loci (33.7% and 35.9% for Cufflinks and StringTie, respectively) were novel, meaning that there were no corresponding gene models in the reference genome annotation. Higher mapping rates were achieved by using the reference genome rather than the *de novo* assembly, with an average of 87.7%, 81.5%, and 69.3% of read pairs being successfully mapped by Tophat. HISAT, and Trinity, respectively (Table 1).

#### Differential expression analysis

All approaches detected changes in expression over time and differences between temperatures (Additional file 1 Table S5). However, the amplitude of the transcriptomic response and, to a lesser extent, the general patterns of differential expression between sample groups differed. For example, the range in differentially expressed genes relative to the baseline is about 5X greater, according to Trinity (0-5121) compared to Cufflinks (147-1272) and StringTie (4-1001). Further, DE increases with temperature at 2 dph, peaks at the intermediate temperature at 14 dph, and is highest at the lower temperature at 29 dph, according to Trinity, whereas it always increases with temperature (StringTie) or varies (Cufflinks) according to the reference-based approaches.

#### Gene ontology enrichment analysis

There was substantial overlap among ClueGO-enriched GO terms associated with differentially expressed genes detected by each pipeline (Figure 1). When more differentially expressed genes were detected by Trinity, the amount of overlap with the reference genome-based pipelines increased, resulting in near concordance between pipelines in those samples that were most different from the baseline (D2-13°C, D14-11°C, and D29-13°C). Conversely, when fewer differentially expressed genes were detected, the discrepancies between the *de novo* and reference-based pipelines were greater. The GO terms identified as being differentially expressed by Cufflinks and StringTie largely, though incompletely, overlapped, with Cufflinks detecting more than StringTie.

**Figure 1:**
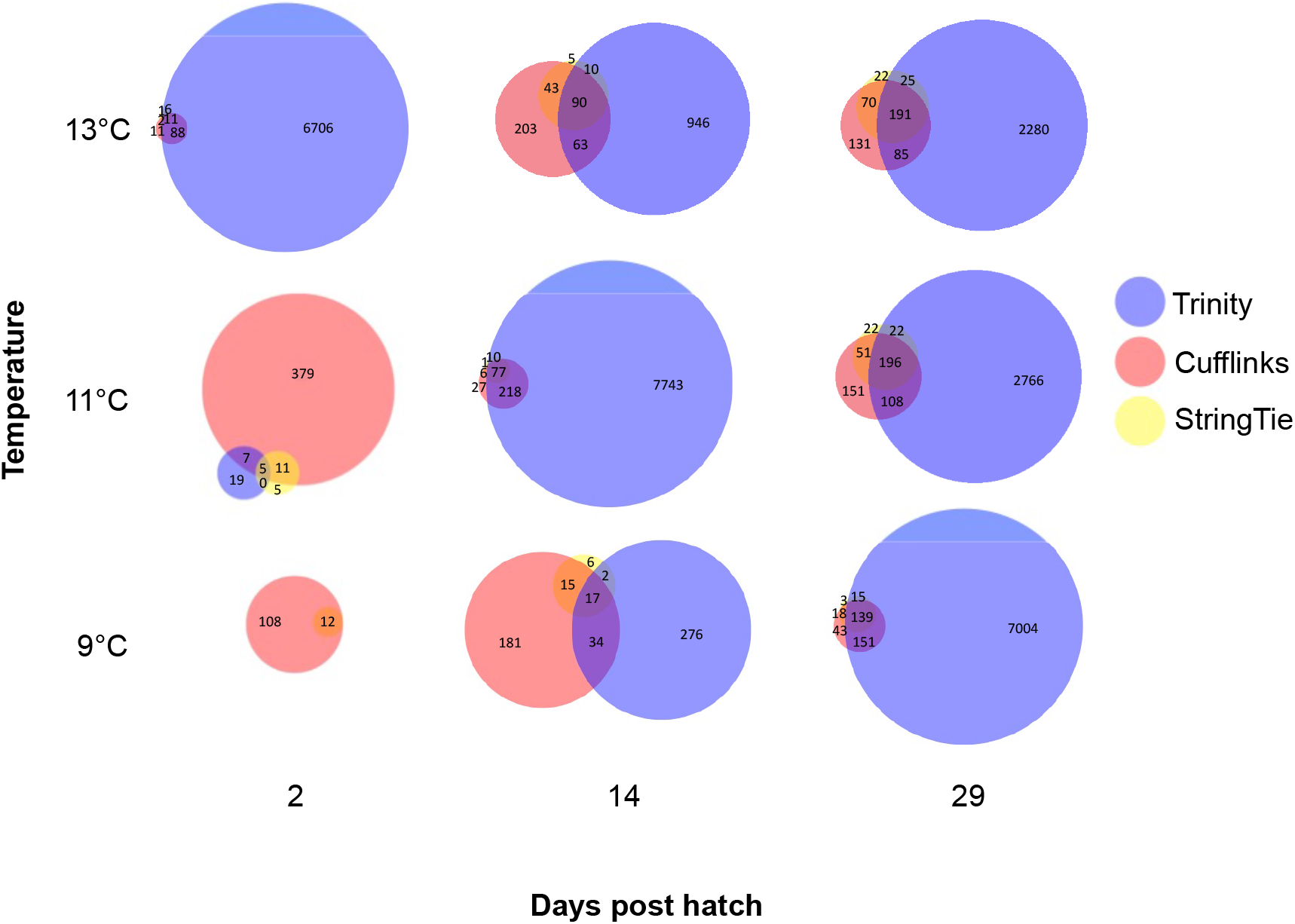
Overlap of gene ontology terms among bioinformatic pipelines. The number of unique gene ontology (GO) terms enriched among differentially expressed genes at each time and temperature in a larval Atlantic cod rearing experiment (Oomen et al. [provisionally accepted at *Integrative & Comparative Biology*]), using the *de novo* Trinity pipeline, the reference-based Tuxedo pipeline (Cufflinks), and the ‘new Tuxedo’ pipeline (StringTie). Enrichment analysis was performed using ClueGO v.2.3.2 [17] in Cytoscape v.3.2.1 [19]. The sizes of circles and overlaps are proportional to the number of GO terms within individual diagrams (i.e., time points × treatments), but not between them.

Notably, however, Trinity also tended to identify more enriched GO terms relative to the number of differentially expressed genes than the other pipelines, with an average of 0.20 (+/upregulated) and 1.33 (-/downregulated) BiNGO-enriched terms per gene for all samples relative to the baseline, compared to 0.07 (+) and 0.06 (−) for Cufflinks and 0.03 (+) and 0.42 (−) for StringTie (Table 2). The same pattern was true for upregulated, but not downregulated, ClueGO-enriched GO terms (Table 2).

**Table 2:**
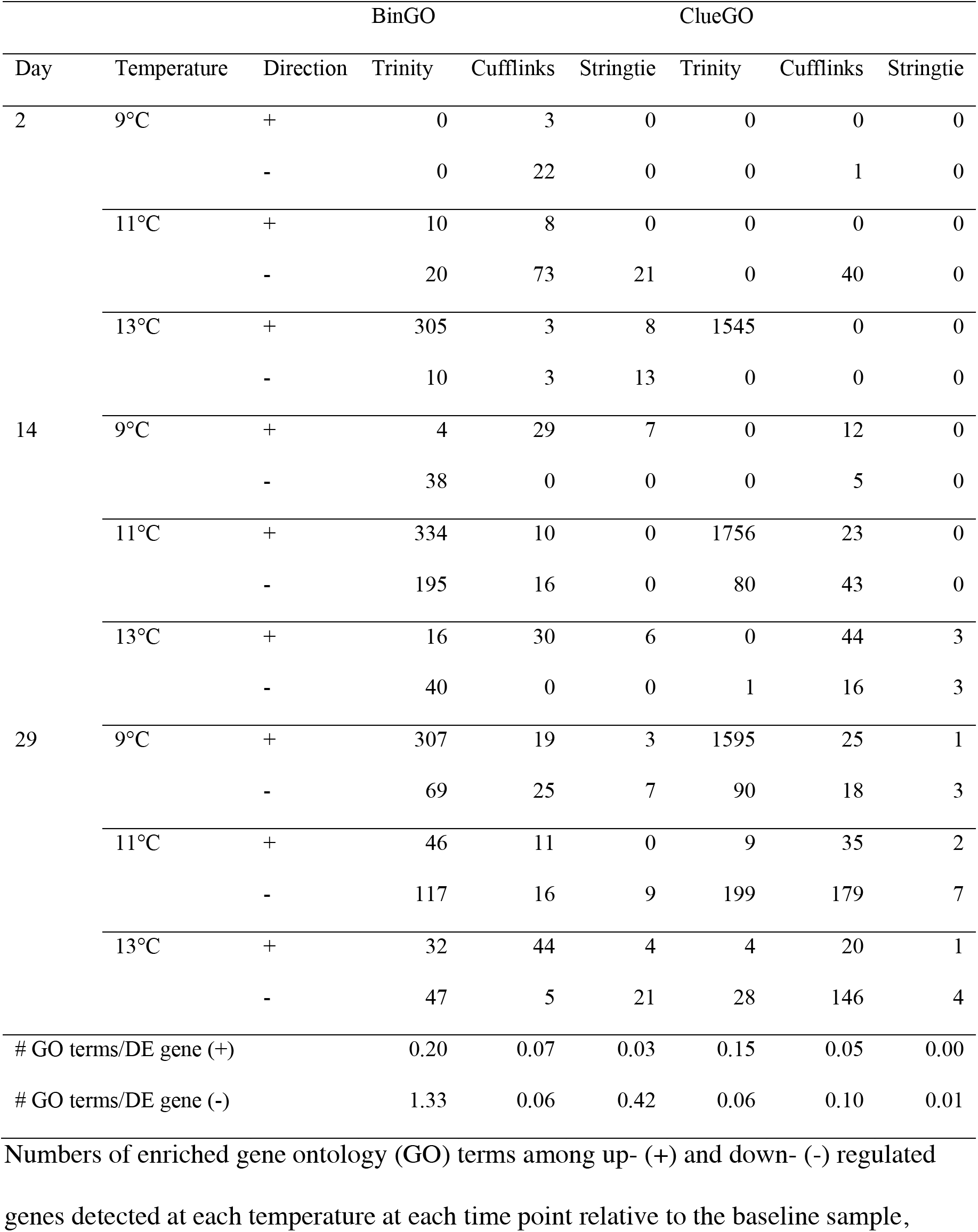
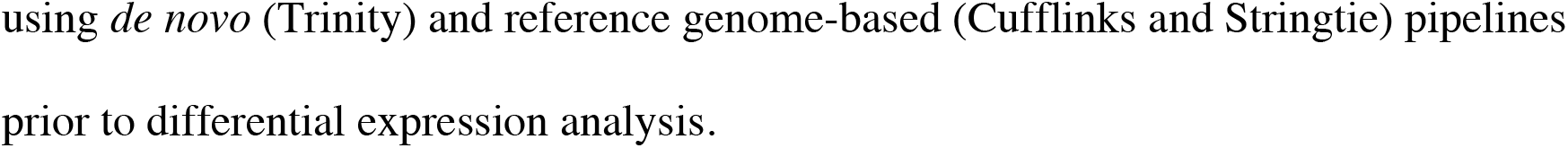
Numbers of enriched gene ontology terms detected through pairwise contrasts with the baseline.

Specifically, the numbers of ClueGO-enriched terms based on upregulated genes detected by Tuxedo/Cufflinks followed a similar pattern as those based on Trinity (despite different trends in the numbers of differentially expressed genes) except with a time lag: no enriched processes were detected at 2 dph, whereas the greatest response was observed at 14 dph at 13°C and 29 dph at 11°C.

### Discussion

Overall, the three pipelines (Trinity, Tuxedo, and the ‘new Tuxedo’) produced different, yet complementary, results. The generally greater number of differentially expressed genes detected by Trinity could be due to *de novo* assembled transcripts that were unable to be mapped to the reference genome. This could be because they were located on genome fragments that were not assembled or filtered out of the reference, or because they were novel isoforms lacking corresponding gene models in the reference [8]. The observation that approximately one-third of all transcripts assembled using the reference genome were novel supports the hypothesis that the reference genome and annotation are incomplete (Additional file 1 Table S4). Indeed, genome assemblies and annotations are ‘works-in-progress’ [21, 22]. Thus, draft genomes of non-model organisms are typically less complete and less accurate than their older, model counterparts. Further, although 10-15% (Cufflinks) and 12-28% (StringTie) of read pairs failed to map to the reference genome (Table 1), these reads might contribute to *de novo* transcripts constructed by Trinity.

The fragmented nature of the *de novo* transcriptome also likely contributed to the greater number of differentially expressed genes detected using Trinity, because fragmentation can result in a single gene being represented by multiple partial transcripts, which are identified as unique ‘genes’ according to Trinity. However, this alone does not explain the large number of unique GO terms relative to the reference genome-based pipelines (Figure 1; Table 2), as different fragments of a gene would have the same ontology.

### Conclusion

Among the reference genome-based approaches, the ‘new Tuxedo’ pipeline [23] outperformed the Tuxedo suite [24]. This was expected, given that StringTie is known for improved reconstruction of low-abundance, multi-isoformic, and highly multi-exonic genes [14]. Greater enrichment by Trinity-identified differentially expressed genes suggests that a high proportion of them represent biologically meaningful differences in transcription, as opposed to transcriptional noise or false positives. Coupled with the ability to annotate novel loci, the increased sensitivity of the *de novo* pipeline for detecting likely biologically meaningful differential expression might make it preferable over the reference genome-based approaches for studies aimed at broadly characterizing variation in the magnitudes of expression differences and biological processes. However, the reference-based ‘new Tuxedo’ pipeline might be more appropriate when a more conservative approach is warranted, such as for the detection and identification of candidate genes.

Overall, we suggest that the comparison and integration of multiple methods is highly informative. We also suggest that the annotation of the Atlantic cod genome could be improved by incorporating our transcriptome data [21, 22].

## Limitations

The present methodological comparison is based on the genome and transcriptome assemblies of a single, non-model species. We do not know how common the discrepancies between pipelines we observed are; direct comparisons between *de novo* and reference genome-based pipelines are rarely reported. Indeed, the differences are likely to vary substantially between species depending on the quality of the genomic resources available.

## List of abbreviations

DE: differential expression
dph: days post hatch
FPKM: Fragments Per Kilobase Mapped
CPM: Counts Per Million
GO: Gene Ontology

## Declarations

### Ethics approval and consent to participate

This study was conducted in accordance with animal welfare laws in Norway and approved by the Animal Ethics Committee run by the Norwegian Food Safety Authority (Mattilsynet).

### Consent for publication

Not applicable.

### Availability of data and material

The RNA-seq dataset analysed in the present study is available at [TBD]. All other data are included in the present paper and Additional File 1.

### Competing interests

The authors declare that they have no competing interests.

### Funding

This research was supported by funding from Canada’s Natural Sciences and Engineering Research Council through a Discovery Grant to JAH and a Canada Graduate Scholarship and Michael Smith Foreign Study Supplement to RAO, the Research Council of Norway to JAH, Interreg IV (MarGen) to HK, a Killam Predoctoral Scholarship and Nova Scotia Graduate Scholarship to RAO, and the Centre for Ecological and Evolutionary Synthesis.

### Authors’ contributions

RAO and JAH designed the study. JAH, HK, and NCS contributed funds. EMO and HK managed wild fish collection and supervised the fish rearing experiment. RAO conducted the fish rearing experiment, molecular laboratory work, and data analyses. SJ managed the RNA sequencing. RAO, SJ, and JAH interpreted the results. RAO wrote the manuscript with feedback from the other authors. All authors read and approved the manuscript.

## Acknowledgements

We acknowledge the contributions of the staff and students at the *Institute of Marine Research, Flødevigen* for their facilities, technical assistance, and additional support during the experiment, especially P. Baardsen, S. Stiansen, R. Johansen, H. Sannaes, K. Halvorsen, and T. Sørdalen. We are grateful to the Centre for Ecological and Evolutionary Synthesis (CEES) for technical and administrative assistance and to M. Solbakken, O.K. Tørresen, and M.H.S. Hansen for enabling the molecular and bioinformatics work. The transcriptome sequencing was carried out at the Norwegian Sequencing Centre (NSC; http://www.sequencing.uio.no), University of Oslo. All computational work was performed on the Abel Supercomputing Cluster (Norwegian metacenter for High Performance Computing (NOTUR) and the University of Oslo) operated by the Research Computing Services group at USIT, the University of Oslo IT-department (http://www.hpc.uio.no/). Thanks to P. Bentzen, C. Herbinger, and M. Johnston for helpful comments and discussion.

## Authors’ information

Not applicable.

## Notes

### Competing Interest Statement

The authors have declared no competing interest.

