## Additional File 1 for "Comparison of *de novo* and reference genome-based transcriptome assembly pipelines for differential expression analysis of RNA sequencing data"

#### Figures

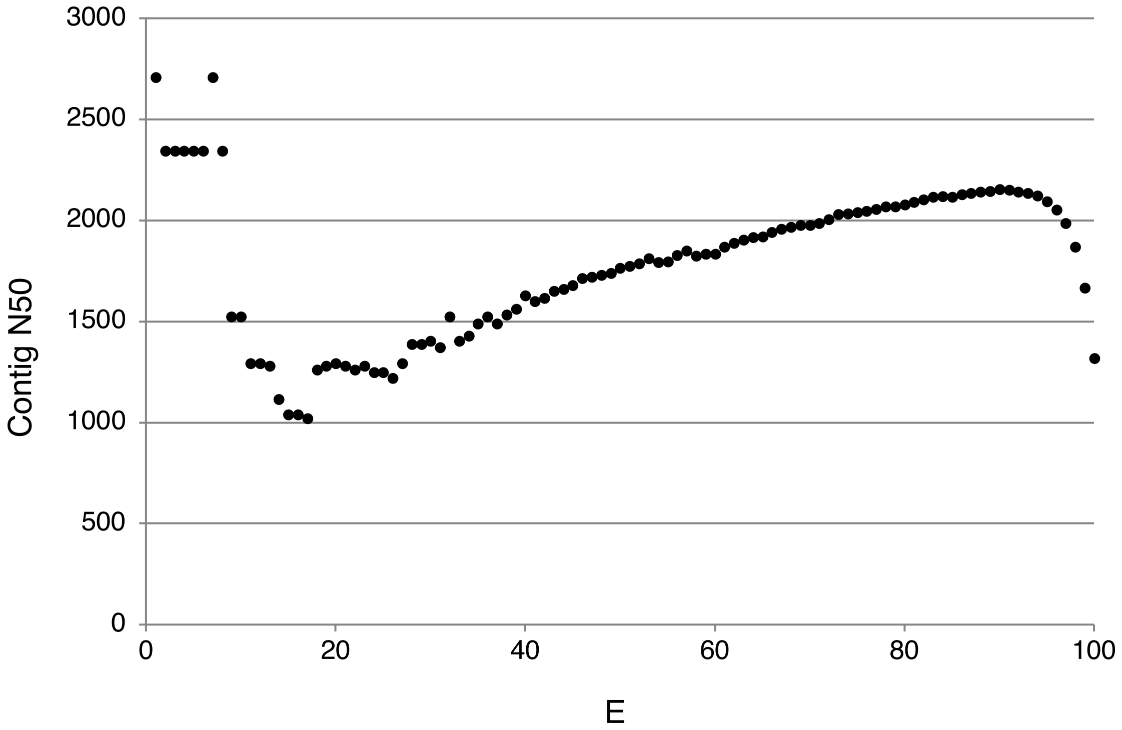


**Figure S1: Expression contig N50 distribution of the *de novo* Trinity assembly.** The minimum length of contig in which 50% of all assembled bases are contained as a function of the percentage of total normalized expression data represented by the most highly expressed trancripts in the *de novo* Trinity assembly.

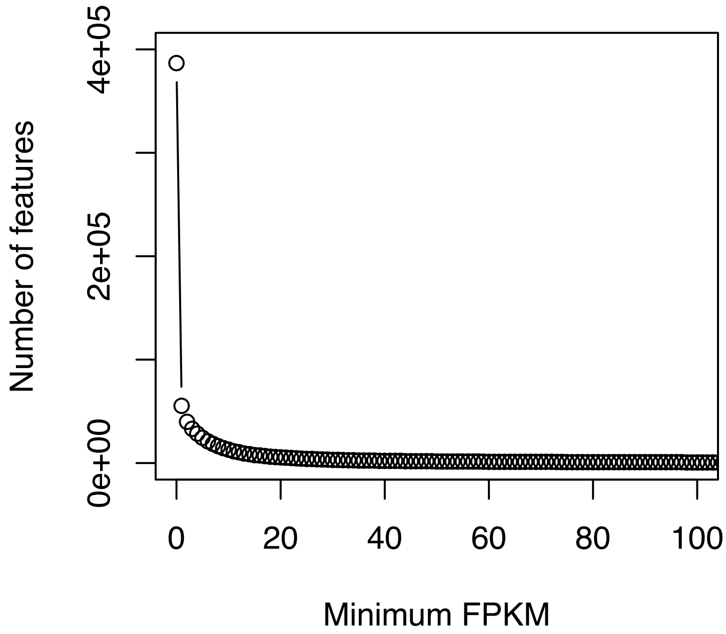


**Figure S2: Minimum expression level gene count.** The number of *de novo*-assembled Trinity 'genes' with a minimum expression level in FPKM (Fragments Per Kilobase of transcript per Million mapped reads).

### Tables

**Table S1: *De novo* transcriptome assembly statistics.**

| Statistic | Value |
|---|---|
| *General* | |
| Total trinity 'genes' | 386872 |
| Total trinity transcripts | 530683 |
| Percent GC | 47.93 |
| *Based on all transcript contigs* | |
| Contig N10 | 3949 |
| Contig N20 | 2841 |
| Contig N30 | 2180 |
| Contig N40 | 1690 |
| Contig N50 | 1278 |
| Median contig length | 431 |
| Average contig | 778.02 |
| Total assembled bases | 412883956 |
| *Based on only the longest isoform per 'gene'* | |
| Contig N10 | 3484 |
| Contig N20 | 2338 |
| Contig N30 | 1648 |
| Contig N40 | 1159 |
| Contig N50 | 835 |
| Median contig length | 364 |
| Average contig | 612.59 |
| Total assembled bases | 236994575 |

**Table S2: *De novo* transcriptome ExN50 statistics.**

| E | E-N50 | # transcripts |
|---|---|---|
| E1 | 2710 | 2 |
| E2 | 2343 | 4 |
| E3 | 2343 | 6 |
| E4 | 2343 | 9 |
| E5 | 2343 | 11 |
| E6 | 2343 | 14 |
| E7 | 2710 | 17 |
| E8 | 2343 | 21 |
| E9 | 1523 | 24 |
| E10 | 1523 | 28 |
| E11 | 1292 | 31 |
| E12 | 1292 | 35 |
| E13 | 1282 | 39 |
| E14 | 1117 | 43 |
| E15 | 1039 | 48 |
| E16 | 1039 | 52 |
| E17 | 1021 | 57 |
| E18 | 1262 | 63 |
| E19 | 1282 | 68 |
| E20 | 1292 | 74 |
| E21 | 1282 | 81 |
| E22 | 1262 | 88 |
| E23 | 1282 | 95 |
| E24 | 1248 | 103 |
| E25 | 1248 | 112 |
| E26 | 1220 | 123 |
| E27 | 1292 | 134 |
| E28 | 1389 | 147 |
| E29 | 1389 | 161 |
| E30 | 1404 | 178 |
| E31 | 1371 | 197 |
| E32 | 1523 | 218 |
| E33 | 1404 | 242 |
| E34 | 1428 | 269 |
| E35 | 1488 | 298 |
| E36 | 1523 | 331 |
| E37 | 1488 | 367 |
| E38 | 1535 | 407 |
| E39 | 1561 | 453 |
| E40 | 1628 | 504 |
| E41 | 1599 | 561 |

| | | |
|---|---|---|
| E42 | 1615 | 625 |
| E43 | 1650 | 698 |
| E44 | 1661 | 780 |
| E45 | 1680 | 872 |
| E46 | 1713 | 973 |
| E47 | 1721 | 1085 |
| E48 | 1730 | 1208 |
| E49 | 1741 | 1343 |
| E50 | 1765 | 1491 |
| E51 | 1775 | 1654 |
| E52 | 1787 | 1833 |
| E53 | 1812 | 2028 |
| E54 | 1794 | 2243 |
| E55 | 1797 | 2477 |
| E56 | 1828 | 2732 |
| E57 | 1851 | 3008 |
| E58 | 1826 | 3304 |
| E59 | 1836 | 3624 |
| E60 | 1836 | 3970 |
| E61 | 1868 | 4340 |
| E62 | 1890 | 4737 |
| E63 | 1905 | 5162 |
| E64 | 1917 | 5619 |
| E65 | 1920 | 6107 |
| E66 | 1943 | 6631 |
| E67 | 1958 | 7191 |
| E68 | 1967 | 7789 |
| E69 | 1976 | 8427 |
| E70 | 1976 | 9107 |
| E71 | 1986 | 9833 |
| E72 | 2007 | 10605 |
| E73 | 2031 | 11426 |
| E74 | 2034 | 12298 |
| E75 | 2042 | 13227 |
| E76 | 2048 | 14219 |
| E77 | 2055 | 15279 |
| E78 | 2068 | 16414 |
| E79 | 2070 | 17631 |
| E80 | 2079 | 18939 |
| E81 | 2092 | 20348 |
| E82 | 2105 | 21871 |
| E83 | 2117 | 23524 |
| E84 | 2119 | 25325 |

| | | |
|---|---|---|
| E85 | 2117 | 27303 |
| E86 | 2129 | 29482 |
| E87 | 2136 | 31915 |
| E88 | 2142 | 34656 |
| E89 | 2144 | 37789 |
| E90 | 2153 | 41413 |
| E91 | 2152 | 45701 |
| E92 | 2143 | 50947 |
| E93 | 2134 | 57639 |
| E94 | 2122 | 66540 |
| E95 | 2095 | 79086 |
| E96 | 2053 | 97771 |
| E97 | 1986 | 127442 |
| E98 | 1871 | 177874 |
| E99 | 1667 | 268326 |
| E100 | 1318 | 488067 |

**Table S3: De novo transcriptome blastx and blastp top hit statistics.**

| Percent length coverage bin | bin count | bin ≥ count |
|---|---|---|
| *blastx* | | |
| 100 | 6888 | 6888 |
| 90 | 2542 | 9430 |
| 80 | 2113 | 11543 |
| 70 | 2034 | 13577 |
| 60 | 2266 | 15843 |
| 50 | 2500 | 18343 |
| 40 | 2627 | 20970 |
| 30 | 2697 | 23667 |
| 20 | 2341 | 26008 |
| 10 | 782 | 26790 |
| *blastp* | | |
| 100 | 7385 | 7385 |
| 90 | 2389 | 9774 |
| 80 | 2046 | 11820 |
| 70 | 1940 | 13760 |
| 60 | 2173 | 15933 |
| 50 | 2304 | 18237 |
| 40 | 2473 | 20710 |
| 30 | 2344 | 23054 |
| 20 | 1763 | 24817 |
| 10 | 432 | 25249 |

**Table S4: Comparison of reference genome-based transcriptome assemblies with the reference genome annotation.**

| Statistic | Value(s) | Value(s) |
|---|---|---|
| | *Cufflinks* | *Stringtie* |
| mRNAs | 167150 | 92033 |
| Loci | 31830 | 34132 |
| Multi-exon transcripts | 158926 | 83848 |
| Multi-transcript loci | 18066 | 17075 |
| Mean transcripts per locus | 5.3 | 2.7 |
| | *Reference* | *Reference* |
| mRNAs | 23243 | 23243 |
| Loci | 23243 | 23243 |
| Multi-exon transcripts | 22139 | 22139 |
| | *Cufflinks vs. Reference* | *Stringtie vs. Reference* |
| Super-loci w/ reference transcripts | 20333 | 20523 |
| Base level sensitivity | 100 | 100 |
| Base level precision | 52.8 | 56.7 |
| Exon level sensitivity | 100 | 100 |
| Exon level precision | 54.6 | 57.4 |
| Intron level sensitivity | 100 | 100 |
| Intron level precision | 61.2 | 64.4 |
| Intron chain level sensitivity | 100 | 100 |
| Intron chain level precision | 13.9 | 26.4 |
| Transcript level sensitivity | 100 | 100 |
| Transcript level precision | 13.9 | 25.3 |
| Locus level sensitivity | 100 | 100 |
| Locus level precision | 65.6 | 62.5 |
| Matching intron chains | 22139 | 22139 |
| Matching transcripts | 23243 | 23243 |
| Matching loci | 23243 | 23243 |
| Missed exons | 0/223454 (0.00%) | 0/223454 (0.00%) |
| Novel exons | 110161/427064 (25.80%) | 94374/415948 (22.70%) |
| Missed introns | 1/200211 (0.00%) | 1/200211 (0.00%) |
| Novel introns | 48063/327303 (14.70%) | 43522/311125 (14.00%) |
| Missed loci | 0/23243 (0.00%) | 0/23243 (0.00%) |
| Novel loci | 10720/31830 (33.70%) | 12252/34132 (35.90%) |
| Total union super-loci across all input datasets | 31830 | 34132 |

**Table S5: Numbers of genes differentially expressed in pairwise tests.** Number of up- (+), down- (-), and dys- (+/-) regulated transcripts detected a) at each temperature at each time point relative to the baseline sample, b) between temperature treatments within time points, and c) between time points within temperature treatments, using de novo (Trinity) and reference genome-based (Cufflinks and Stringtie) pipelines prior to differential expression analysis.

| Day | Temperature | Direction | Trinity | Cufflinks | Stringtie |
|---|---|---|---|---|---|
| *a) All vs. baseline (0 dph at 9°C)* | | | | | |
| 2 | 9°C | + | 0 | 49 | 2 |
| | | - | 0 | 98 | 2 |
| | | +/- | 0 | 147 | 4 |
| | 11°C | + | 8 | 97 | 29 |
| | | - | 2 | 452 | 8 |
| | | +/- | 10 | 549 | 37 |
| | 13°C | + | 3581 | 70 | 67 |
| | | - | 65 | 110 | 13 |
| | | +/- | 3646 | 180 | 80 |
| 14 | 9°C | + | 48 | 177 | 64 |
| | | - | 46 | 179 | 41 |
| | | +/- | 94 | 356 | 105 |
| | 11°C | + | 4296 | 274 | 114 |
| | | - | 825 | 278 | 61 |
| | | +/- | 5121 | 552 | 175 |
| | 13°C | + | 188 | 392 | 279 |
| | | - | 106 | 372 | 110 |
| | | +/- | 294 | 764 | 389 |
| 29 | 9°C | + | 3558 | 366 | 318 |
| | | - | 718 | 352 | 205 |
| | | +/- | 4276 | 718 | 523 |
| | 11°C | + | 509 | 465 | 499 |
| | | - | 720 | 607 | 377 |
| | | +/- | 1229 | 1072 | 876 |
| | 13°C | + | 629 | 429 | 677 |
| | | - | 439 | 630 | 324 |
| | | +/- | 1068 | 1059 | 1001 |
| *b) Temperature effects* | | | | | |
| 2 | 9-11°C | + | 0 | 16 | 5 |
| | | - | 0 | 32 | 1 |
| | | +/- | 0 | 48 | 6 |
| | 11-13°C | + | 3291 | 114 | 7 |

| | | | | | |
|---|---|---|---|---|---|
| | | - | 64 | 65 | 10 |
| | | +/- | 3355 | 179 | 17 |
| | 9-13°C | + | 3556 | 70 | 29 |
| | | - | 49 | 78 | 8 |
| | | +/- | 3605 | 148 | 37 |
| 14 | 9-11°C | + | 1507 | 38 | 4 |
| | | - | 30 | 18 | 2 |
| | | +/- | 1537 | 56 | 6 |
| | 11-13°C | + | 156 | 90 | 24 |
| | | - | 2324 | 92 | 11 |
| | | +/- | 2480 | 182 | 35 |
| | 9-13°C | + | 4 | 103 | 19 |
| | | - | 0 | 47 | 5 |
| | | +/- | 4 | 150 | 24 |
| 29 | 9-11°C | + | 0 | 80 | 13 |
| | | - | 0 | 66 | 18 |
| | | +/- | 0 | 146 | 31 |
| | 11-13°C | + | 1 | 26 | 24 |
| | | - | 0 | 59 | 8 |
| | | +/- | 1 | 85 | 32 |
| | 9-13°C | + | 0 | 46 | 11 |
| | | - | 0 | 55 | 6 |
| | | +/- | 0 | 101 | 17 |
| *c) Time effects* | | | | | |
| 9°C | 0*-2 | + | 0 | 49 | 2 |
| | | - | 0 | 98 | 2 |
| | | +/- | 0 | 147 | 4 |
| | 2-14 | + | 50 | 470 | 33 |
| | | - | 36 | 349 | 13 |
| | | +/- | 86 | 819 | 46 |
| | 14-29 | + | 206 | 166 | 81 |
| | | - | 118 | 168 | 27 |
| | | +/- | 324 | 334 | 108 |
| | 2-29 | + | 3663 | 388 | 229 |
| | | - | 835 | 387 | 109 |
| | | +/- | 4498 | 775 | 338 |
| 11°C | 0*-2 | + | 8 | 97 | 29 |
| | | - | 2 | 452 | 8 |
| | | +/- | 10 | 549 | 37 |

| | | | | | |
|---|---|---|---|---|---|
| | 2-14 | + | 4580 | 835 | 72 |
| | | - | 2447 | 485 | 70 |
| | | +/- | 7027 | 1320 | 142 |
| | 14-29 | + | 546 | 219 | 86 |
| | | - | 2946 | 218 | 43 |
| | | +/- | 3492 | 437 | 129 |
| | 2-29 | + | 1392 | 1261 | 816 |
| | | - | 2958 | 1222 | 1319 |
| | | +/- | 4350 | 2483 | 2135 |
| 13°C | 0*-2 | + | 3581 | 70 | 67 |
| | | - | 65 | 110 | 13 |
| | | +/- | 3646 | 180 | 80 |
| | 2-14 | + | 167 | 456 | 134 |
| | | - | 804 | 320 | 103 |
| | | +/- | 971 | 776 | 237 |
| | 14-29 | + | 22 | 102 | 42 |
| | | - | 10 | 219 | 20 |
| | | +/- | 32 | 321 | 62 |
| | 2-29 | + | 375 | 399 | 417 |
| | | - | 978 | 496 | 370 |
| | | +/- | 1353 | 895 | 787 |
| # of transcripts in analysis: | | | 51075 | 31830 | 34064 |
